# A NanoDam toolkit for tissue-specific transcription factor profiling in *C. elegans*

**DOI:** 10.1101/2023.05.31.543105

**Authors:** Callista Yee, Yutong Xiao, Dimitris Katsanos, Taylor N. Medwig-Kinney, Wan Zhang, Kang Shen, David Q. Matus, Michalis Barkoulas

**Affiliations:** Howard Hughes Medical Institute, Stanford University, Stanford, CA 94305, USA; Department of Biology, Stanford University, Stanford, CA 94305, USA; Department of Biochemistry and Cell Biology, Stony Brook University, Stony Brook, NY 11794, USA; Department of Life Sciences, Imperial College, London SW7 2AZ, UK; Wu Tsai Neurosciences Institute, Stanford University, Stanford, CA 94305, USA

**Author notes:** Correspondence to Callista Yee and Michalis Barkoulas. These authors contributed equally (Callista Yee, Yutong Xiao). Present Address: Department of Biology, University of North Carolina at Chapel Hill, Chapel Hill, NC 27599, USA. Arcadia Science, Berkeley, CA 94704, USA.

**Keywords:** C. elegans, genomics, DamID, RNT-1/RUNX, EGL-43/EVI1, Anchor Cell, Neurons

## Abstract

During development of multicellular organisms, cells must execute precise molecular decisions to achieve cell fate specification and differentiation. These decisions are orchestrated by networks of transcription factors (TFs) which act to regulate gene expression of specific cohorts of genes to ultimately confer identity. Depending on the cellular context, TF expression can vary dramatically both spatially and temporally. These differences in expression patterns can result in tissue-specific differences in TF binding to downstream targets. To identify targets on a tissue-specific basis, Targeted DamID (TaDa) has been recently introduced to generate TF binding profiles in various models including *C. elegans*. However, TaDa suffers from portability such that a new promoter-TF fusion transgene must be constructed for every new experimental condition of interest. Here, we adapt NanoDam for usage in *C. elegans*, which relies on endogenous TF-GFP knock-ins, a plethora of which have already been generated by the community. We report that NanoDam single copy transgenes consisting of lowly expressed, tissue-specific GFP nanobody-Dam fusions, when combined with endogenous GFP-tagged alleles of TFs, results in robust, tissue-specific profiling. Using an endogenous GFP-tagged allele of EGL-43/EVI1, we performed NanoDam profiling of two disparate tissue types, the anchor cell (AC) and dopaminergic neurons, and identify targets unique to each and shared by both cell types. We also identify two GATA TFs, ELT-6 and EGL-18, as novel regulators of AC invasion. Together, we demonstrate that NanoDam is capable of profiling endogenous GFP-tagged TFs to identify novel downstream targets in specific cell types of *C. elegans*.

## Introduction

During development, transcription factors (TFs) play critical roles in executing gene expression programs that provide cellular identity and specification. For example, through differential expression of TFs, sister cells may possess dramatically different terminal morphology and function (PACKER *et al*. 2019). Decades of research have identified TFs involved in all aspects of *C. elegans* development, but their genetic targets, on a cell-type specific basis, remain relatively underexplored (REINKE *et al*. 2013). Endogenous labelling experiments have revealed that some TFs are expressed in diverse tissue types and that their temporal expression can vary; specific isoforms of TFs can be expressed during short windows of time in one cell type, while others may be continuously expressed in others until death (SHERWOOD *et al*. 2005; AHN *et al*. 2022; MASOUDI *et al*. 2023).

<u>Ch</u>romatin Immuno<u>p</u>recipitation (ChIP)-seq has been widely regarded as the gold standard technique to determine DNA sequences that are bound by TFs of interest (PARK 2009). The majority of ChIP-seq studies in *C. elegans* have been generated by the modENCODE consortium and utilize whole animal lysates expressing exogenous transgenes(REINKE *et al*. 2013). While these datasets represent a valuable resource, they do have limitations as, first, whole worm lysates may mask cell-specific biology, and second, transgenes may not be fully representative of the endogenous TF expression thereby introducing artificial TF binding events. To this end, proximity-based labelling using Dam methylase has become a common alternative (GREIL *et al*. 2006). A tissue-specific version of this technique, TaDa (Targeted DamID), has been successfully utilized in various models including *C. elegans* to identify genomic loci that may be bound by TFs of interest (SOUTHALL *et al*. 2013; KATSANOS *et al*. 2021; KATSANOS AND BARKOULAS 2022).

To perform this DamID-based profiling, a single copy TaDa transgene is inserted into the genome at a defined chromosomal site. TaDa transgenes are comprised of a tissue-specific promoter driving a TF::Dam methylase fusion (SOUTHALL *et al*. 2013) (Figure 1A). Upon binding of the TF to DNA, Dam will permanently methylate DNA in a proximity-based fashion, where it will add a methyl group to the N6 of adenine at the recognition sequence 5’-GATC-3’ (Figure 1A). Total DNA is subsequently harvested at the developmental stage of interest, and methylated DNA is selectively digested, amplified, and sequenced to determine TF binding. Although this technique has been reliably used to generate tissue-specific profiling information, it requires a new Dam-TF fusion insertion to be generated for every tissue-specific promoter and TF of interest, which can be laborious to streamline.

**Figure 1:**
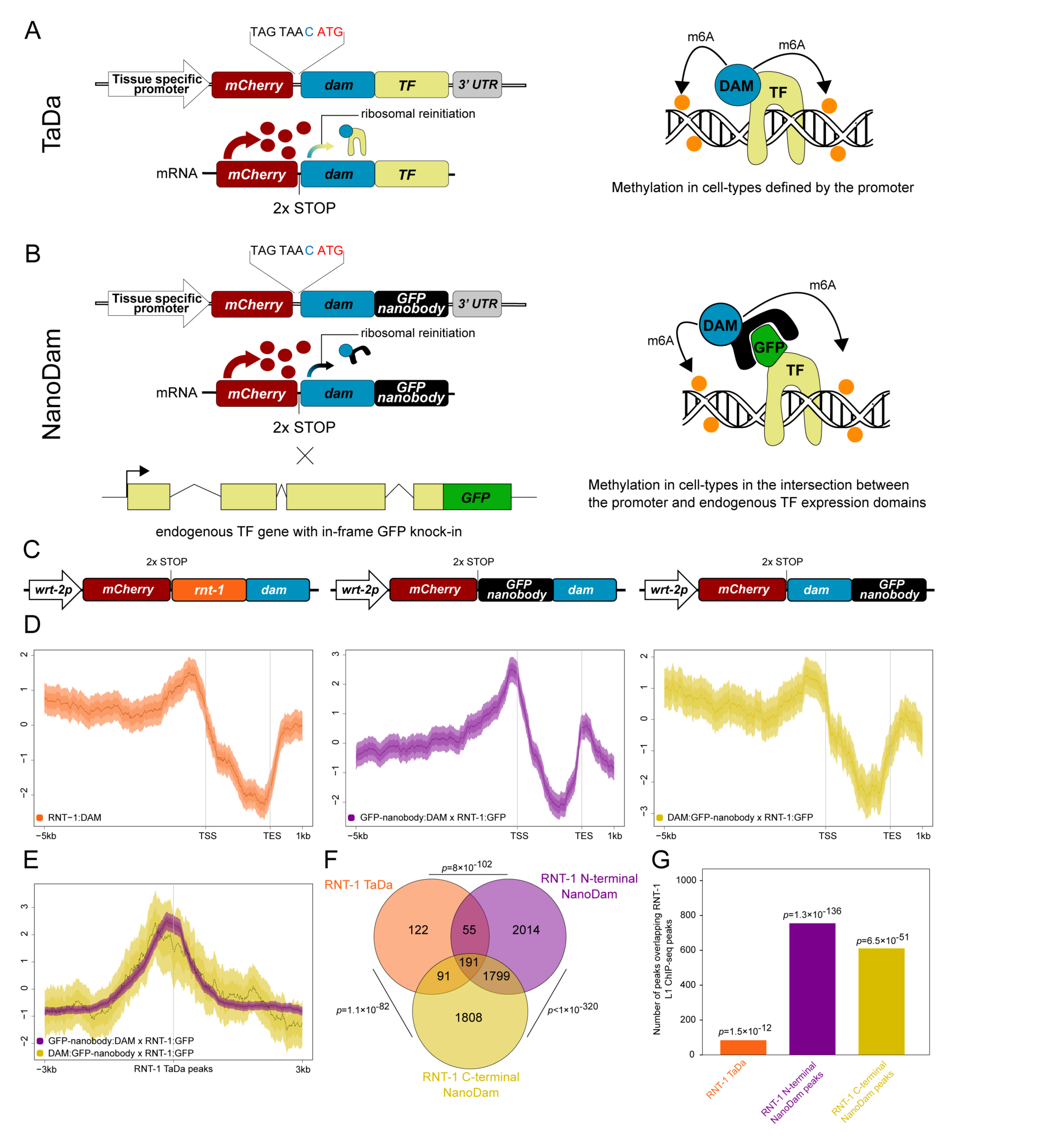
The NanoDam system exceeds performance of the TaDa system to generate TF binding profiles. A) Overview of the TaDa and NanoDam systems. Left: Schematic of TaDa transgenes. A tissue specific promoter is used to drive bicistronic expression of two ORFs. The first ORF encodes a red fluorophore (mCherry), followed by two stop codons and an extra nucleotide for frameshift. The second ORF encodes Dam methylase fused to a TF of interest. The resulting mRNA will encode both ORFs, but the second ORF will be translated at low levels due to ribosomal reinitiation. Right: Model depicting tissue-specific methylation action of Dam::TF fusions. Dam methylase will methylate the N6 adenine of the consensus sequence 5’-GATC-3’-in a proximity dependent fashion. B) Overview of the NanoDam system. Left: Schematic of NanoDam transgenes. NanoDam transgenes mirror TaDa transgenes but utilize a Dam::GFP nanobody fusion instead of a direct Dam::TF fusion. NanoDam transgenes are then crossed into animals carrying endogenous GFP-tagged TFs to allow for tissue-specific TF profiling. Right: Model depicting tissue-specific methylation action of Dam::GFP-nanobody fusions. C) Design of TaDa and NanoDam transgenes to generate RNT-1 binding profiles in the seam cells for comparisons across methods. Left: RNT-1 TaDa. Center: N-terminal NanoDam. Right: C-terminal NanoDam. D) Aggregation plots of profiling data generated by RNT-1 TaDa and NanoDam. Average enrichment scores (z-scores on y-axis) are shown in 10-bp bins for regions of equal length across all specified genomic features (x-axis). Aggregation plots are drawn to label features of genes, including 5-kb upstream of the transcription start site (TSS), 1-kb downstream of the transcription end site (TES), and gene bodies condensed into a 2-kb pseudo-length. Shaded areas represent 95% confidence intervals. E) Aggregation plot comparing C- and N-terminal NanoDam signal against all RNT-1 TaDa peaks. A window of 3-kb upstream and 3-kb downstream of RNT-1 TaDa peaks is shown to highlight signal intensity of NanoDam datasets is centered at TaDa peaks. F) Venn diagram plots of all called peaks between RNT-1 TaDa and C- and N-terminal NanoDam. *P* values are calculated by Monte Carlo simulations for significance of overlaps between datasets. G) Bar graph depicting number of peak overlaps between RNT-1 TaDa and NanoDam profiling and RNT-1 whole animal ChIP-seq at the L1 stage.

To circumvent this demand, a system coined NanoDam (Nanobody – DamID) has been recently described in *Drosophila melanogaster* (TANG *et al*. 2022). In this method, instead of a direct TF::Dam fusion, a vhh4 GFP nanobody is fused to Dam methylase to generate tissue-specific NanoDam driver lines (Figure 1B). NanoDam driver lines can then be crossed to animals carrying endogenous GFP-tagged TFs to acquire tissue-specific TF profiling (Figure 1B). This bipartite system is particularly advantageous within model organism communities as it allows to take advantage of the large collection of readily available GFP-tagged TF alleles generated by the community that can be used in combination with suitable driver lines.

Here, we sought to adapt the NanoDam system and generate the first toolkit in *C. elegans.* First, we compared the efficacy of NanoDam to the widely adopted Targeted DamID (TaDa) system by generating data for the same TF and using the same tissue-specific promoter. Second, we applied NanoDam to study the binding of the same GFP-tagged TF to different targets in different tissues, the somatic gonad-derived anchor cell (AC) and a subset of sensory neurons. Last, we validate binding events identified by NanoDam and identify new TFs that play a role in the process of AC invasion, a critical morphogenetic process associated with uterine-vulval development. This tool will enable the *C. elegans* community to perform rapid tissue-specific, low input genomic profiling of TFs in a streamlined fashion.

## Materials and Methods

### *C. elegans* strains and culture conditions

Animals were cultured under standard conditions and grown at 20–25°C on NGM plates seeded with *E. coli* OP50 as previously described (BRENNER 1974). For all NanoDam experiments, gravid adult worms were treated with alkaline hypochlorite to isolate eggs (PORTA-DE-LA-RIVA *et al*. 2012). Eggs were resuspended in M9 buffer and allowed to hatch overnight at room temperature to obtain synchronized L1s. Synchronized L1s were placed on NGM plates seeded with *E. coli dam-/-dcm-/-* (NEB strain ER2925) and grown for at least 3 generations without starvation. Worms were subsequently grown in bulk, synchronized, and then harvested at the stage of interest.

### Molecular biology

Plasmids used in this study were generated using either standard restriction digest cloning or Gibson assembly techniques. NanoDam cassettes (mKate2::Dam::VHH4, mCherry::Dam::VHH4 and mCherry::VHH4::Dam) were generated using synthetic DNAs from either IDT or Twist Biosciences. Schematics for the NanoDam cassettes are depicted in Figure 1B.

To construct the conventional TaDa RNT-1:Dam fusion plasmid driven by the epidermal promoter of the *wrt-2* gene, the oligos DK150 and DK151 were used to amplify the *rnt-1* coding sequence from N2 cDNA. The amplicon carrying compatible overhangs was inserted by Gibson assembly in a XmaJI digested pPB7 vector (KATSANOS AND BARKOULAS 2022) generating pDK93(*wrt-2p::mCherry::rnt-1:dam::unc-54 3’UTR + cb-unc-119*). NanoDam plasmids made up of the same components were built for direct comparisons against TaDa in capturing RNT-1 binding. To this end, the *egl-13 NLS::vhhGFP4* cassette was amplified from pYX013 using oligo pairs DK269 and DK270 as well as DK14 and PB25. The resulting amplicons were inserted by Gibson assembly into either an XmaJI or a PaeI digested pDK7 vector respectively to generate pDK172(*attR4-L1::mCherry::egl-13NLS:vhhGFP4:linker:Dam-myc::unc-54 3’UTR + cb-unc-119*) and pDK173(*attR4-L1::mCherry::Dam:linker:egl-13NLS:vhhGFP4::unc-54 3’UTR + cb-unc-119*). The promoter of *wrt-2* was cloned upstream of the NanoDam fusions utilising the above two plasmids and pKA279 (Ward et al., 2013 PLOS Genetics) to generate pDK175(*wrt-2p::mCherry::egl-13NLS:vhhGFP4:linker:Dam-myc::unc-54 3’UTR + cb-unc-119*) and pDK179(*wrt-2p::mCherry::Dam:linker:egl-13NLS:vhhGFP4::unc-54 3’UTR + cb-unc-119*). All pDK plasmids generated in this study were constructed using the pCFJ151 vector backbone and contain homology to a safe harbor site corresponding to the MosSCI site ttTi5605.

To build the remaining NanoDam driver plasmids, tissue specific promoters and NanoDam cassettes were amplified and cloned using Gibson Assembly into pYX013, which contains a C-terminal NanoDam cassette and a selectable self-excising cassette (SEC) with homology to a safe harbor corresponding to the MosSCI site ttTi4348. The promoter of *lin-29* was cloned using oligo pairs DQM521 and DQM522 to generate pYX8 (*SEC::lin-29p::mKate2:STOP:Dam:linker:egl-13NLS:vhh4GFP::unc-54 3’UTR)*. The promoter of *dat-1* was cloned from pCM560 using oligo pairs DQM1113 and DQM1114 to generate pCY35 (*SEC::dat-1p::mKate2:STOP:Dam:linker:egl-13NLS:vhh4GFP::unc-54 3’UTR)*. The promoter of *rgef-1* was cloned from pSYL62 using oligo pairs DQM1138 and DQM119 to generate pCY38 (*SEC::rgef-1p::mKate2:STOP:Dam:linker:egl-13NLS:vhh4GFP::unc-54 3’UTR)*. The promoter of *ser-2* was cloned from pCM316 using oligo pairs CY693 and CY694 to generate pCY39 (*SEC::ser-2prom3::mKate2:STOP:Dam:linker:egl-13NLS:vhh4GFP::unc-54 3’UTR)*.

### Transgenesis

Single copy insertions of NanoDam drivers were generated by *Mos1*-mediated single-copy insertion (MosSCI) (FROKJAER-JENSEN *et al*. 2014) or by the SEC method using CRISPR/Cas9 genome editing via standard microinjection into the hermaphrodite gonad (DICKINSON *et al*. 2015). Hermaphrodite adults were co-injected with guide plasmid (50 ng/μl), repair plasmid (50 ng/μl) and an extrachromosomal array marker (pCFJ90, 2.5 ng/μl). Injected animals and their progeny were incubated at 25°C for three days before carrying out genotyping to identify successfully edited loci. Following identification of CRISPR-mediated insertions, floxing of the SEC cassette was carried out using standardized protocols (DICKINSON *et al*. 2015).

To generate GFP, NLS-GFP or mNeonGreen endogenously tagged alleles of transcription factors, DNA encoding fluorophores was amplified using PCR with primers providing at least 50 bp of homology to the target locus. Plasmids pCM552 (GFP and NLS-GFP) and pJW2171 (mNeonGreen) were used as templates. Amplified PCR products were pooled together and purified using a Qiagen PCR Purification kit. Concentrated PCR products were eluted in water and measured using a NanoDrop instrument (ThermoFisher Scientific). 0.5 µL of 100 µM cRNA (IDT) targeting either the C- or N-terminal locus of interest was added to 3 µL of 10 µM tracrRNA (IDT) and heated to 95°C for 5 minutes and cooled to 25°C for 5 minutes. The resulting cRNA:tracrRNA reaction was added to a 1.5 mL tube containing 0.5 µL of ALT-R Cas9 (IDT) and heated in a 37°C water bath for 15 minutes. During the incubation, the purified fluorophore PCR amplicon was diluted to 300 ng/µL and melted in a PCR machine (GHANTA AND MELLO 2020). To complete the injection mixture, 1 µL of co-injection plasmid (pRF4, final concentration 50 ng/µL) and 5 µL of melted PCR product was added to the 1.5 mL tube. Mixes were spun at 13000 RPM for 15 minutes and kept on ice during microinjection.

### Extraction and amplification of methylated DNA

Synchronized animals carrying NanoDam transgenes with or without the GFP-tagged TF of interest were cultured on *E. coli dam-/-dcm-/-* bacteria and grown to the appropriate stage of interest. Animals were washed 3x in M9 buffer and snap frozen in liquid nitrogen and stored at −80°C until needed as previously described (KATSANOS *et al*. 2021). At least three biological replicates were collected for each condition (NanoDam transgene alone; NanoDam transgene + GFP allele) and all samples were processed together to avoid batch effects. In the case of the RNT-1 TaDa and NanoDam comparison experiments 2 biological replicates were grown simultaneously and were collected, washed and frozen as previously described (KATSANOS AND BARKOULAS 2022). gDNA and methylated DNA was purified using a Qiagen DNeasy blood and tissue kit and processed as previously described (GOMEZ-SALDIVAR *et al*. 2021). All subsequent steps used to purify and amplify methylated DNA by NanoDam are identical to standard *C. elegans* TaDa protocols. gDNA was digested using DpnI (New England Biolabs) to cut methylated DNA and double strand adaptors were ligated using T4 DNA Ligase (ThermoFisher Scientific). Freshly ligated DNA was amplified using HiDi DNA Polymerase (myPols Biotech) and analyzed on a 1% gel to assess amplification. PCR was performed using 4 initial long cycles, followed by 14-20 additional cycles as previously described (KATSANOS *et al*. 2021). PCR products were purified using magnetic beads (Beckman) or a QIAquick PCR Purification Kit (Qiagen) and quantified using either a Nanodrop or a Qubit 4.0 and a dsDNA High Sensitivity kit (ThermoFisher Scientific).

### Sequencing

Sequencing was performed using either Illumina or Nanopore platforms. For Illumina sequencing, library preparation and next-generation sequencing was performed by GENEWIZ on a HiSeq4000 instrument. For Nanopore sequencing, libraries were prepared according to manufacturer protocols using a Ligation Sequencing Kit (SQK-LSK-109, ONT) and a Native Barcoding Expansion Kit (EXP-NBD104 and EXP-NBD114, ONT). Sequencing of pooled libraries was performed on a Nanopore MinION Mk1C instrument using Spot-ON flow cells (FLO-MIN106D). Basecalling and demultiplexing was performed onboard the MinION Mk1C using fast basecalling with a minimum Q quality score of 8.

### Calculation of TaDa and NanoDam signal profiles and peak calling

The quality of all resulting FASTQ files were validated using FastQ stats (available from https://github.com/owenjm/damid_misc/blob/master/fastq-stats). Data pertaining to RNT-1 profiling experiments were sequenced using the Illumina platform and the resulting fastq files were used as input to the damidseq_pipeline (v.1.5.3) (MARSHALL AND BRAND 2015) using bowtie2 (v.2.3.4.1) to map reads to the *C. elegans* WBcel235 genome assembly. Generated bam files from each TF or control strain for TaDa or NanoDam were used to perform all pairwise calculation between control and TF replicates to produce bedGraph DamID signal files as previously described (KATSANOS AND BARKOULAS 2022). Signal across replicates was averaged to a single profile by quantile normalisation and peak calling was performed using MACS2 with subsequent statistical thresholding and peak-merging as described (TANG *et al*. 2022). Peaks were assigned to nearby genes using UROPA (KONDILI *et al*. 2017).

Data pertaining to EGL-43 profiling experiments were sequenced using the Nanopore platform and were similarly mapped using minimap2 (v. 2.26) to the *C. elegans* ce10 genome and GATC bins. Samtools (v.1.17) was used to manipulate alignment files to generated sorted .BAMs, which were fed into the damidseq_pipeline (v.1.5.3) (MARSHALL AND BRAND 2015). Biological experiments were paired together with either a free Dam control or a NanoDam only control replicate. Binding profiles for each TF and condition was then generated by averaging the binding intensities across all comparisons per GATC-bin. Averaged intensities were transformed as previously described (TANG *et al*. 2022) and peaks were called from bedGraph files (generated from Dam/control pairwise comparisons) using the find_peaks software (freely available from https://github.com/owenjm/find_peaks)). Peaks were filtered with an FDR < 0.01 and overlapping peaks were merged using bedtools (v.2.26.0). Visualization of peaks was performed using IGV (v.2.15.2). Peaks were assigned to the closest transcriptional start site using bedtools (v.2.26.0) and annotataed using HOMER (v.4.11). Aggregation plots were generated using SeqPlots (STEMPOR AND AHRINGER 2016). Plots represent signal averages in 10 bp bins around specified positional features flanking genes, as previously described (KATSANOS *et al*. 2021).

### Calculation of overlaps between gene profiling experiments

Overlaps between gene profiling experiments were calculated using bedtools in a pairwise fashion. For RNT-1 comparison experiments overlaps and their significance was calculated by Monte Carlo simulations using OLOGRAM (FERRE *et al*. 2021). For the rest, statistical significance of overlaps between experiments was calculated using a Fisher’s exact test using bedtools as previously described (KATSANOS *et al*. 2021). Venn diagrams were generated using either Intervene (KHAN AND MATHELIER 2017) or http://bioinformatics.psb.ugent.be/webtools/Venn/ or GraphPad Prism (v.9).

### Peak enrichment analysis

Identification of enriched gene ontology (GO) terms or association with tissue specific expression for the genes associated with peaks identified in this study was performed using PANTHER (biological process, Fisher overrepresentation test, FDR correction) (ASHBURNER *et al*. 2000; GENE ONTOLOGY 2021) and the Tissue Enrichment tool available on Wormbase (*q*-value threshold = 0.1) (ANGELES-ALBORES *et al*. 2016) respectively.

### ChIP-qPCR

Worms were bleach synchronized and grown at 25°C until they reached the L3 stage, then washed 3X in PBS containing Complete Mini protease inhibitor (Roche). Worms were briefly spun down, resuspended in a minimal amount of PBS and dripped into liquid nitrogen to create worm popcorn. Worm popcorn was kept at −80°C until 3 biological replicates were harvested. Popcorn was ground in a mortar and pestle in liquid nitrogen until a fine powder. Next, 10 volumes of PBS (containing 1.1% formaldehyde, Complete Mini protease inhibitor (Roche) and phosphatase inhibitor cocktail (Calbiochem)) was added to the powder and incubated for 10 minutes at RT with gentle rocking. Formaldehyde was quenched by adding glycine to a final concentration of 125 mM and the reaction was pelleted and washed in ice cold PBS containing 1 mM PMSF. The resulting pellet was resuspended in buffer FA (50mM HEPES/KOH pH 7.5, 1 mM EDTA, 1% Triton X-100, 0.1% sodium deoxycholate, 150mM NaCl) supplemented with 0.1% sarkosyl and protease and phosphatase inhibitors. The resuspended pellet was then sonicated using an EpiShear probe sonicator (Active Motif) for 2×13 minutes at 80% power (30 sec on, 30 sec off). Once sonication was complete, IP was performed using GFP-Trap Agarose (Chromotek) according to manufacturer protocols. Unconjugated agarose beads were used as a mock control. 25 uL of the reaction was reserved to extract input DNA. IP and input DNA were purified using ChIP elution buffer (250 mM NaCl, 1% SDS, 10 mM Tris-Cl pH 8, 1mM EDTA) and incubation with proteinase K (ThermoFisher Scientific) overnight at 65°C. Both IP and input DNA was purified using a PCR Purification Kit (Qiagen). DNA was quantified using a Qubit 4.0 with a dsDNA High Sensitivity kit (ThermoFisher Scientific). qPCR using primers targeting the peaks of interest was performed on the resulting purified input and IP DNA using SsoFast EvaGreen SuperMix on a Bio-Rad CFX96 thermocycler (Bio-Rad).

### Auxin-induced protein degradation and RNA interference (RNAi)

For all auxin experiments, synchronized L1 animals were cultured on regular NGM plates until the P6.p 2-cell stage (mid-L3 stage) before being subsequently transferred to NGM plates containing either 0.1mM 5-Ph-IAA or 4mM K-NAA. 5-Ph-IAA or K-NAA solutions were freshly prepared and diluted in molten NGM agar (cooled to 50°C) immediately before pouring plates as previously described (MARTINEZ AND MATUS 2020). Plates were seeded using overnight cultures of OP50. OP50-seeded NGM plates supplemented with 0.25% ethanol were utilized for control conditions. Single copy TIR1 transgenes *wrdSi22* (*eft-3p::TIR1::P2A::BFP-AID\**) and bmd284 (loxN, rpl-28p::TIR1(F79G)::T2A::DHB::2xmKate2) were used to perform auxin-induced degradation.

RNAi against *egl-43, egl-18* and *elt-6* was performed on synchronized L1s using feeding vectors as previously described (MEDWIG-KINNEY *et al*. 2020). Synthetic DNAs corresponding to *egl-18* and *elt-6* were generated by Twist Biosciences as DNA fragments and subsequently cloned into restriction-digested T444T using NEB Gibson Assembly to generate RNAi feeding vectors.

### Imaging of NanoDam and endogenous fluorophore tagged TFs

Animals were mounted into a drop of M9 on a 5% agarose pad containing 10 mM sodium azide or 10 mM levamisole and covered with a glass coverslip. All imaging was performed either on an upright Zeiss AxioImager A2 with a Borealis-modified CSU10 Yokogawa spinning disk scan head (Nobska Imaging) using 440, 514, 488, and 561 nm Vortran lasers in a VersaLase merge and a Plan-Apochromat 100×/1.4 (NA) or 40×/1.4NA Oil DIC objective with a Hamamatsu Orca EM-CCD camera controlled by MetaMorph, or an inverted Zeiss Axio Observer Z1 microscope equipped with a Yokogawa CSU-W1 spinning-disk unit using 488 and 561 nm lasers and a Prime 95B Scientific CMOS camera with a C-Apochromat 40×0.9 NA objective controlled by 3i Slidebook software.

### Assessment of AC invasion and specification

The invasion of the AC into the vulval epithelium was scored by the presence of a gap in the basement membrane (BM), situated underneath the AC. A LAM-2::mNG transgene was used to label the basement membrane and assess AC invasion. In the wild type, successful invasion is defined as a breach as wide as the basolateral surface of the AC at the P6.p 4-cell stage (SHERWOOD AND STERNBERG 2003).

### Image quantification and analysis

All images were processed and analysed using ImageJ/Fiji (v.2.9) and the SUM stack function. Regions of interest corresponding to the nuclei of neurons and the anchor cell were circled manually in ImageJ and Mean Grey Values were recorded. Detailed quantification of nuclei intensity and anchor cell invasion was performed as previously described (MEDWIG-KINNEY *et al*. 2020; XIAO *et al*. 2023). Statistical significance was calculated using two-tailed Student’s *t*-tests (GraphPad Prism (v.9)).

## Results

### NanoDam design and comparison to TaDa

To generate NanoDam driver strains, we followed a similar design as previously described(TANG *et al*. 2022) and fused the nanobody vhhGFP4 to either the N- or C-terminus of Dam methylase. We utilized well characterized promoters to drive expression of the NanoDam system as a single copy insertion (Table 1, Table S1). Similar to the design of TaDa (Figure 1A), NanoDam relies on the production of a bicistronic message (Figure 1B) as follows. First, a tissue-specific promoter drives expression of a primary open reading frame which contains the coding sequence of a red fluorophore (to allow visualization of tissue-specific expression), followed by two stop codons (TAG and TAA) and a single-nucleotide frameshift. Following this, a secondary open reading frame contains the coding sequence of the nanobody-Dam fusion protein. This bicistronic message is generated through ribosome re-initiation and allows for expression of the secondary ORF at low levels. This low expression level of the nanobody-Dam fusion is critical to prevent saturation of methylation levels and methylation induced toxicity (KATSANOS AND BARKOULAS 2022). Once NanoDam transgenes are generated by single-copy insertion into the genome, they can be crossed together with animals expressing endogenous GFP-tagged TFs (Figure 1B).

**Table 1:** List of NanoDam Driver Strains.

| Strain | Allele | Promoter | Dam Fusion to Nanobody | Tissue |
| --- | --- | --- | --- | --- |
| DQM986 | *bmdSi245* | *lin-29* | C-terminal | Anchor Cell |
| DQM954 | *bmdSi261* | *dat-1* | C-terminal | Dopaminergic Neurons |
| DQM1041 | *bmdSi282* | *rgef-1* | C-terminal | Pan-neuronal |
| DQM1294 | *bmdSi364* | *ser-2* (prom 3) | C-terminal | PVD, PDE and OLL |
| MBA1362 | *icbSi228* | *wrt-2* | C-terminal | Seam cells/epidermis |
| MBA1360 | *icbSi226* | *wrt-2* | N-terminal | Seam Cells/epidermis |

To test the ability of NanoDam to generate accurate profiling information, we compared the conventional TaDa system to both C-terminal and N-terminal nanobody::Dam fusions. We utilized an endogenous GFP-tagged allele of *rnt-1*, the sole *C. elegans* ortholog of human RUNX transcription factors. RNT-1 is expressed in many cells, including the *C. elegans* epidermis where it functions together with its binding partner BRO-1/CBF-Beta to promote symmetric cell divisions of the seam cells (KAGOSHIMA *et al*. 2005; NIMMO *et al*. 2005; KATSANOS AND BARKOULAS 2022). The availability of an endogenous GFP CRISPR knock-in for RNT-1, and our interest in beginning to understand the mechanisms via which it promotes seam cell identity, motivated us to use it as a platform to compare the performance of TaDa and NanoDam for the same TF.

To assess if NanoDam can capture RNT-1 binding in a similar manner to TaDa, we generated single copy insertions of *wrt-2p::rnt-1::dam* (TaDa)*, wrt-2p::vhh4::dam* (C-terminal NanoDam) and *wrt-2p::dam::vhh4* (N-terminal NanoDam)(Figure 1C). To capture the downstream targets of RNT-1, we performed profiling of RNT-1 TaDa and RNT-1 NanoDam (C-and N-terminal) at the L4 stage. We identified 459 statistically significant peaks through TaDa and 3894 and 4063 peaks through C- and N-terminal NanoDam (complete list in Table S2). Encouragingly, as expected for TF binding, we found that all three conditions yielded enrichment of methylation signal primarily upstream and proximal to the transcription start site (TSS) of genes, as visualized through genome-wide aggregation plots (Figure 1D).

Next, we compared the overlap between the RNT-1 NanoDam with RNT-1 TaDa both at the level of signal as well as peak overlaps. We observed significant preference of C- and N-terminal RNT-1 NanoDam signal across RNT-1 peaks identified by TaDa, as opposed to adjacent genomic regions (3kb upstream and downstream) (Figure 1E), thus, indicating that RNT-1 NanoDam is very likely capturing similar binding preference for RNT-1 as TaDa. Despite the substantially fewer statistically significant peaks identified from our RNT-1 TaDa dataset, we found that more than half of them overlapped with either C- or N-terminal RNT-1 NanoDam peaks (191/459 peaks common across all 3 datasets, Figure 1F). Comparisons across the peaks obtained from the C- and N-terminal NanoDam fusions showed approximately a 50% overlap between datasets (1990/3889 for N/C (51%) and 1990/4059 for C/N (49%)).

Having demonstrated a high degree of concordance across our TaDa and NanoDam datasets, we aimed to assess how well each technique captures RNT-1 binding based on independent ChIP-seq experiments. We compared our TaDa and NanoDam profiling data to an existing whole-animal L1-staged ChIP-seq dataset generated by the ENCODE consortium, hosted on the modERN database (Accession number: ENCSR597WQR). This RNT-1 ChIP-seq experiment identified 5560 unique peaks. We found 84 peaks that overlapped between RNT-1 TaDa and ChIP-seq in comparison to 755 and 611 peaks overlapping between N- or C-terminal NanoDam and ChIP-seq RNT-1 respectively (Figure 1G). Although the ChIP-seq experiment was performed at L1 and TaDa and NanoDam were performed at L4, the methylation experiments capture the whole developmental history of RNT-1 binding, which could in part explain some differences in overlap. Taken together, these results demonstrate that both C- or N-terminal NanoDam is largely consistent with TaDa and can capture TF binding in the seam cells.

### Using NanoDam to study targets of the same transcription factor in two different tissues

Next, we wanted to test the ability of NanoDam to identify cell-specific binding profiles in two disparate tissues that both express the same TF. We turned to the proto-oncogene EGL-43/MECOM, a conserved C2H2 zinc finger TF which has been studied extensively in the context of uterine and vulval development (HWANG *et al*. 2007; RIMANN AND HAJNAL 2007; DENG *et al*. 2020; MEDWIG-KINNEY *et al*. 2020). Mutation or RNAi knockdown of *egl-43* leads to cell cycle and cell invasion/migration phenotypes in key cells that contribute to the morphogenesis of the hermaphrodite vulva (MATUS *et al*. 2010; DENG *et al*. 2020; MEDWIG-KINNEY *et al*. 2020). In addition to its role in gonad development, EGL-43 has also been shown to play roles in intestinal development and neuronal migration (GARRIGA *et al*. 1993; RASMUSSEN *et al*. 2013) raising the question how can a single TF play such diverse roles in different tissues? At the L3 stage, EGL-43 is highly expressed in the invasive anchor cell (AC) where it facilitates a pro-invasion program, which is critical for forming the connection between the uterine and vulval tissues so that adult animals can passage eggs to the external environment (SHERWOOD AND STERNBERG 2003). We analyzed a previously reported ChIP-seq dataset of L3 worms carrying endogenously tagged EGL-43 (DENG *et al*. 2020), but we did not observe significant enrichment of migratory or cytoskeletal associated GO terms (Table S3).

To demonstrate the utility of NanoDam, we aimed to dissect the roles of EGL-43 in different tissues using the same GFP-tagged allele. We generated NanoDam drivers expressed specifically in the invasive AC (*lin-29p*) or in dopaminergic (DA) neurons (*dat-1p*) (Figure 2A-B) and crossed each of the drivers into animals carrying endogenously tagged EGL-43::GFP. We performed NanoDam profiling on synchronized L3 animals and harvested 3 independent biological replicates, grown on different days, for each paired condition (NanoDam driver + EGL-43::GFP vs. NanoDam driver alone). All samples were processed in parallel to eliminate batch effects.

**Figure 2:**
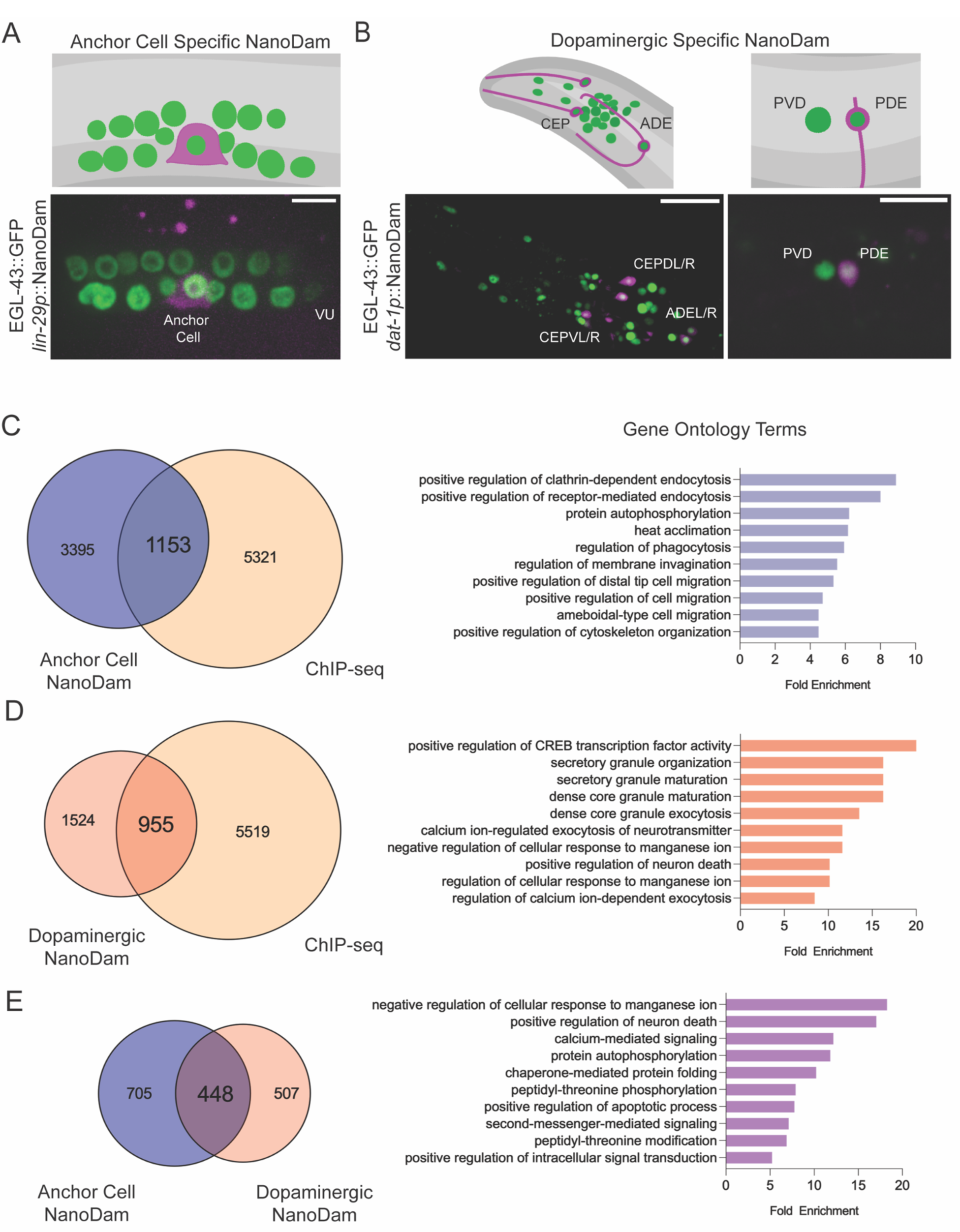
NanoDam profiling of endogenous EGL-43 binding in two distinct tissues. A) Schematic of AC-specific NanoDam. A NanoDam cassette driven using the *lin-29* promoter (*Plin29::mKate2::2xSTOP::vhh4::dam)* is expressed solely in the AC (labelled in magenta) during the L3 stage. EGL-43 (labelled in green), is expressed both in the anchor cell and ventral uterine tissue. Top: cartoon schematic of labeling strategy. Bottom: confocal micrographs of EGL-43::GFP and *lin-29p::*NanoDam. Scale bar = 10μm. Schematic of dopaminergic neuron (DA) specific NanoDam. A NanoDam cassette driven using the *dat-1* promoter (*Pdat-1::mKate2::2xSTOP::vhh4::dam)* is expressed in 3 types of DA neurons (ADE, CEP and PDE). EGL-43 (labelled in green) is expressed in numerous sensory neurons, including all 3 types of DA neurons. Top: schematic of labeling strategy. Bottom: confocal micrographs of EGL-43::GFP and *dat-1p*::NanoDam. Scale bar = 10μm. C) Comparison of peaks identified by EGL-43 anchor cell-specific NanoDam profiling and EGL-43 whole-animal ChIP-seq. Left: Venn diagram representation of identified peaks. Right: Overlapping peaks were assigned identity to nearby genes and subjected to PANTHER GO term analysis. Representative gene ontology terms are plotted with respect to fold enrichment. D) Comparison of peaks identify by EGL-43 DA-specific NanoDam profiling and EGL-43 whole animal ChIP-seq. Left: Venn diagram representation of identified peaks. Right: PANTHER GO term analysis, similar to C). E) Comparison of the overlapping peaks found in the unions of C) and D) to identify common downstream mechanisms of EGL-43. Left: Venn diagram representation. Right: PANTHER GO term analysis, similar to C) and D).

EGL-43 NanoDam profiling in the AC revealed a significant overlap when compared to whole-animal ChIP-seq (1153/6474, Fisher’s exact test *p =* 1.9141e-75, Figure 2C). Tissue-enrichment analyses for the overlapping genes revealed the AC as a top scoring ontology term (*p* < 1.7e-06) with a 3.3-fold enrichment in genes associated with AC expression (Table S4). In comparison, tissue enrichment analysis on genes associated with whole-animal ChIP-seq peaks yielded the AC as a low scoring ontology term (*p* = 0.029) with a 1.3-fold enrichment of genes associated with AC expression. This lesser enrichment might be attributed to the fact that *egl-43* is expressed in many cells, including multiple sensory neurons, contributing to lack of specificity in the dataset.

We found that both AC-specific NanoDam and whole-animal ChIP-seq datasets possessed peaks associated with genes known to genetically function downstream of EGL-43, including the lamelipodin homolog, *mig-10* and the fat-like protocadherin, *cdh-3* (Figure 3A-B). Critically, the AC-specific NanoDam dataset, but not the ChIP-seq dataset, identified binding to some genes that have been reported to be directly regulated by EGL-43, including members of the DNA licensing/replication machinery, *cdt-2,* and *mcm-7* (Figure 3C-D) (DENG *et al*. 2020). Although we detected some binding to these genes in the ChIP-seq dataset, they were not called as statistically significant peaks. A recent study describing the transcriptome of AC reported that *tct-1*, the sole TCTP homolog, regulates matrix metalloproteinases and plays a key role during AC invasion (COSTA *et al*. 2023). We detected EGL-43 binding peaks upstream of *tct-1* through NanoDam, but not ChIP-seq, indicating that EGL-43 may specifically regulate *tct-1* in the AC (Figure 3E). To assess this profiling dataset on a larger scale, we performed GO term analysis and observed many biological terms associated with endocytosis, migration and growth (Figure 2C), supporting the known role of EGL-43 in the AC.

**Figure 3:**
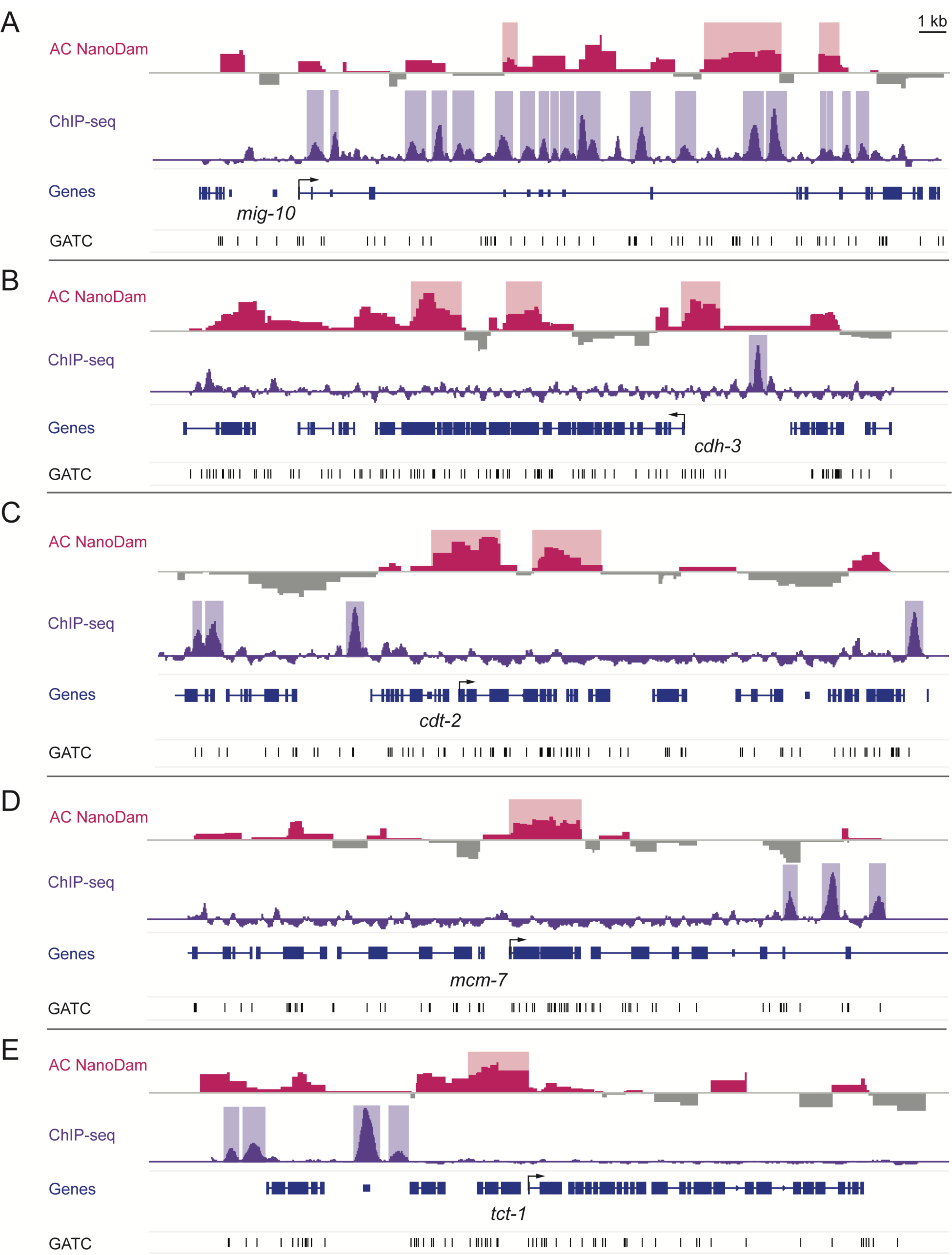
EGL-43 NanoDam and ChIP-seq binding profiles at loci associated with AC development. For each locus, binding profiles for EGL-43 AC NanoDam (red) and EGL-43 ChIP-seq (purple) are presented relative to genomic position (blue) and GATC sites. Computed peaks are denoted by lightly shaded rectangles on the NanoDam and ChIP-seq tracks. A) the *mig-10* locus on chr:III B) the *cdh-3* locus on chr:III C) the *cdt-2* locus on chr:V D) the *mcm-7* locus on chr:V and E) the *tct-1* locus on chr:I.

In a similar fashion, we observed a significant overlap between EGL-43 whole animal ChIP-seq and DA-specific EGL-43 NanoDam (955/6474 peaks, Fisher’s exact test *p* = 1.0304e-157) (Figure 2D). To date, the role of EGL-43 in neurons has been only examined in the context of neuroblast migration and phasmid neuron development. In line with these previously described observations, our GO term analysis resulted in slight enrichment for terms associated with actin filaments and migration (Table S3). However, we found that the most enriched GO terms were associated with secretory granules, neurotransmitter release and endocytosis (Figure 2D). This result was unexpected as EGL-43 has never been described to play roles in any of these processes. Our profiling experiments of EGL-43 in the DA neurons thus support its known role in neuronal migration but also reveal an unexplored potential role in neuronal function.

We compared the overlaps of the ChIP-seq dataset and our DA and AC specific NanoDam experiments to see if EGL-43 is regulating similar downstream targets in two disparate cell types. We observed 448 overlapping peaks, corresponding to a 39% and 47% overlap between datasets (Figure 2E). GO term analysis resulted in terms involved in phosphorylation and intracellular signalling, highlighting kinases such as the serine/threonine nemo-like kinase *lit-1* and the calcium/calmodulin dependent kinases *unc-43* and *ckk-1*. LIT-1 has been well described for its role in establishing polarity signals and has been previously shown to function genetically downstream of EGL-43 in the AC (MATUS *et al*. 2010). It is tempting to speculate that EGL-43 may also function in DA neurons to establish neuronal polarity, as many of the mechanisms used to drive polarity are used by both neurons and the AC (e.g. UNC-6/UNC-40 (ZIEL *et al*. 2009)).

Next, we analyzed the peaks that were unique to the DA and AC NanoDam datasets to identify novel cell-specific functions of EGL-43. We observed 507 and 705 unique peaks, corresponding to 61% and 53% of the respective datasets (Figure 2E). Tissue-enrichment analysis of the peaks unique to the AC were strongly associated to all vulval precursor cells (P3.p-P8.p) and the anchor cell (Table S4), with no apparent enrichment of neuronal cells. GO term analysis of the AC-unique peaks revealed enrichment for terms associated with vulval fate specification and receptor signalling, highlighting the EGFR homolog *let-23* and the RAS homolog *let-60* (Table S3). Similarly, analysis of the peaks unique to DA neurons revealed tissue enrichment of many sensory neurons, including all DA neurons (Table S3). GO term analysis of the DA-unique peaks resulted in enrichment of terms associated with neurotransmitter release and synaptic function, featuring genes such as the synaptotagmin homologs *snt-3* and *snt-7* and the protein kinase C homolog *pkc-1* (Table S3). Taken together, these results demonstrate that NanoDam can provide distinct cell-specific targets of transcription factors which may not be readily identified through traditional whole-animal ChIP-seq profiling.

### NanoDam target validation helps revise known interactions in gene regulatory networks

To validate our NanoDam dataset, we performed ChIP-qPCR on DNA bound to endogenous EGL-43::GFP at the L3 stage. We targeted the peaks corresponding to promoter regions of known downstream genetic targets of EGL-43, such as the cyclin dependent kinase inhibitor *cki-1,* the Nemo-like kinase *lit-1,* and the lamellipodin homolog *mig-10*, as well as the promoter of a novel target, the GATA transcription factor *elt-6* (Figure S1). Additionally, we targeted a peak corresponding to an uncharacterized regulatory region in the *egl-43* locus which was present in both NanoDam datasets (Figure 4A). We found that all putative targets exhibited significant binding to EGL-43 compared to the control (Figure 4B). Notably, we observed that there was approximately a 100-fold enrichment of EGL-43 bound to the uncharacterized regulatory region of the *egl-43* locus (Figure 4B).

**Figure 4:**
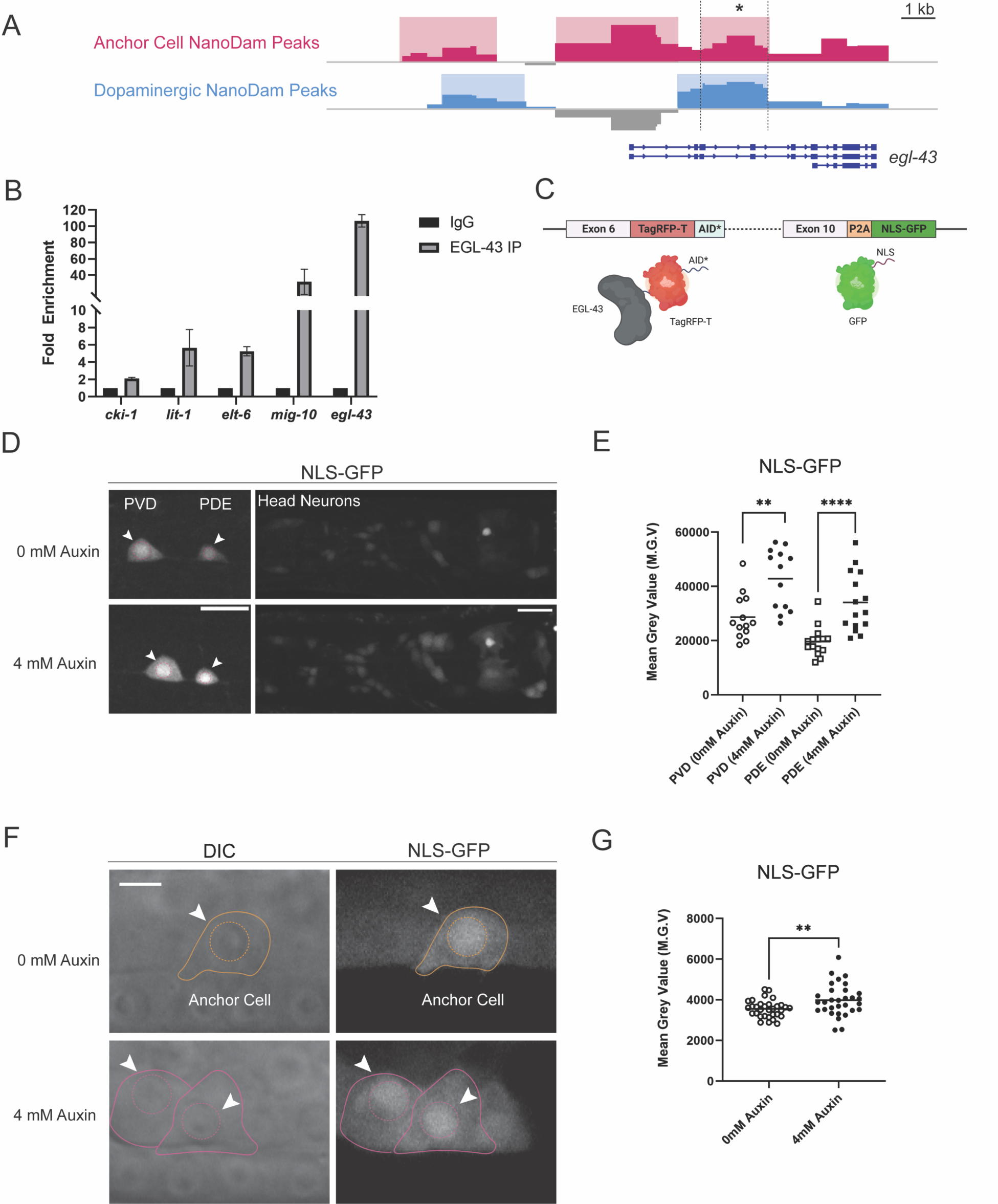
Autoregulation of *egl-43* and ChIP-qPCR validation. A) Schematic of the *egl-43* locus and NanoDam peaks. *egl-43* encodes two isoforms of EGL-43, a long and short isoform. EGL-43 NanoDam peaks mapping to the *egl-43* locus are represented in red (AC-specific NanoDam) and blue (DA neuron-specific NanoDam). A peak which is common to both NanoDam datasets is flanked by dashed lines and an asterisk. B) ChIP-qPCR on endogenous EGL-43 targeting peaks (Figure S1) that were identified using NanoDam. Black bars represent an IgG control; grey bars represent EGL-43 IP (anti-GFP). C) Design of a dual transcriptional and translational endogenous reporter of *egl-43, egl-43(wy1735)*. A single *egl-43* mRNA will generate EGL-43::TagRFP-T::AID::EGL-43 protein and an NLS-GFP due to the introduction of a P2A self-cleaving peptide. D) Representative micrographs of neurons in animals carrying *wrdSi22* and *egl-43(wy1735)*. White arrows denote PVD and PDE and magenta circles outline nuclei. Top: animals treated with 0mM K-NAA auxin; bottom: animals treated with 4mM K-NAA auxin. Scale bar = 10μm E) Quantification of NLS-GFP signal in PVD and PDE nuclei shown in D). F) Confocal micrographs of the anchor cell in animals carrying *wrdSi22* and *egl-43(wy1735)*. White arrows denote AC nuclei and cell outlines are labelled in orange and magenta tracing. Top: animals treated with 0mM K-NAA auxin; bottom: animals treated with 4mM K-NAA auxin. Scale bar = 10μm. G) Quantification of NLS-GFP signal in the AC in animals shown in F). **** denotes a *p*-value of < 0.001 and ** denotes a *p-*value of < 0.01 using a Student’s *t-*test. Scale bar = 5μm.

We next wanted to understand the putative autoregulatory function of *egl-43* and if this behavior is consistent between tissues. The *egl-43* locus encodes two isoforms, a short and long isoform, which have been reported to function redundantly in the AC(DENG *et al*. 2020). NanoDam profiling of EGL-43 in both the AC and DA neurons yielded peaks positioned 5’ to the first exon of both isoforms. Previous work describing the autoregulatory role of *egl-43* demonstrated that *egl-43(RNAi)* resulted in a reduction of intensity of a *Pegl-43::GFP* transcriptional reporter in the AC, indicating positive regulation (MATUS *et al*. 2010). However, this work utilized solely the promoter of the long isoform to drive reporter expression and did not include the additional putative regulatory region that we observed.

To test all potential native autoregulatory elements, we inserted a P2A::NLS-GFP at the C-terminus (exon 10) of the *egl-43* loci in a strain already carrying TagRFP-T::AID* fused to exon 6. This edit resulted in a strain which provides dual transcriptional and translational readout of EGL-43, as well as the ability to degrade EGL-43 in a spatial-temporal fashion. The endogenous NLS-GFP localized highly to nuclei and its expression was consistent with the known expression pattern of EGL-43. Auxin-induced degradation of EGL-43 led to a dramatic increase in nuclear expression of the endogenous NLS-GFP transcriptional reporter (Figure 4D). We quantified intensity of the nuclei of PVD and PDE sensory neurons and observed a significant increase in GFP intensity of 49% and 73% respectively (Figure 4E). Similarly, we observed that loss of EGL-43 resulted in a modest 12% increase in NLS::GFP signal in the AC, consistent with our results that we observed in neurons (Figure 4F). Although our observations are inconsistent with the previous reports of positive autoregulation of *egl-43*, those experiments excluded a putative key regulatory sequence in the reporter transgene (MATUS *et al*. 2010; MEDWIG-KINNEY *et al*. 2020). This result suggests that in both neurons and ACs, EGL-43 is likely to directly negatively regulate its own expression. It also demonstrates that NanoDam is a useful tool to identify novel regulatory regions, thereby enriching our understanding of gene regulatory networks.

### Using NanoDam to expand gene regulatory networks

Next, we explored our dataset to see if we could identify new TF players in AC invasion that are perhaps directly regulated by EGL-43. Interestingly, we identified two GATA TFs, *egl-18* and *elt-6*, which are paralogs and are physically located adjacent to each other on the genome. In the somatic gonad, we found that both TFs were expressed highly in the AC and at low levels in the neighboring ventral uterine cells (VU), an expression pattern that is strikingly reminiscent of the NR2E1 nuclear hormone receptor TF, NHR-67 (Figure 4A). NHR-67 has been described to function downstream of EGL-43 during invasion (FERNANDES AND STERNBERG 2007; MEDWIG-KINNEY *et al*. 2020), raising the possibility that EGL-18/ELT-6 may also be regulated by EGL-43 in a similar manner.

First, we performed RNAi knockdown of *egl-43* and examined the expression of endogenous mNeonGreen reporters of EGL-18 and ELT-6. We found that *egl-43* RNAi led to a dramatic reduction of EGL-18 and ELT-6 in the AC, but not in adjacent VUs, resulting in expression levels that were similar to those normally found in the VU cells (Figure 5A-B). To further confirm this result, we performed auxin-induced degradation of EGL-43 and found that expression of endogenous EGL-18 and ELT-6 in the AC was similarly abolished (Figure 5C-D). Together with our ChIP-qPCR experiment validating binding of EGL-43 to the *elt-6* locus (Figure 4B), these results provide evidence that EGL-43 may directly regulate expression of these two TFs in the AC.

**Figure 5:**
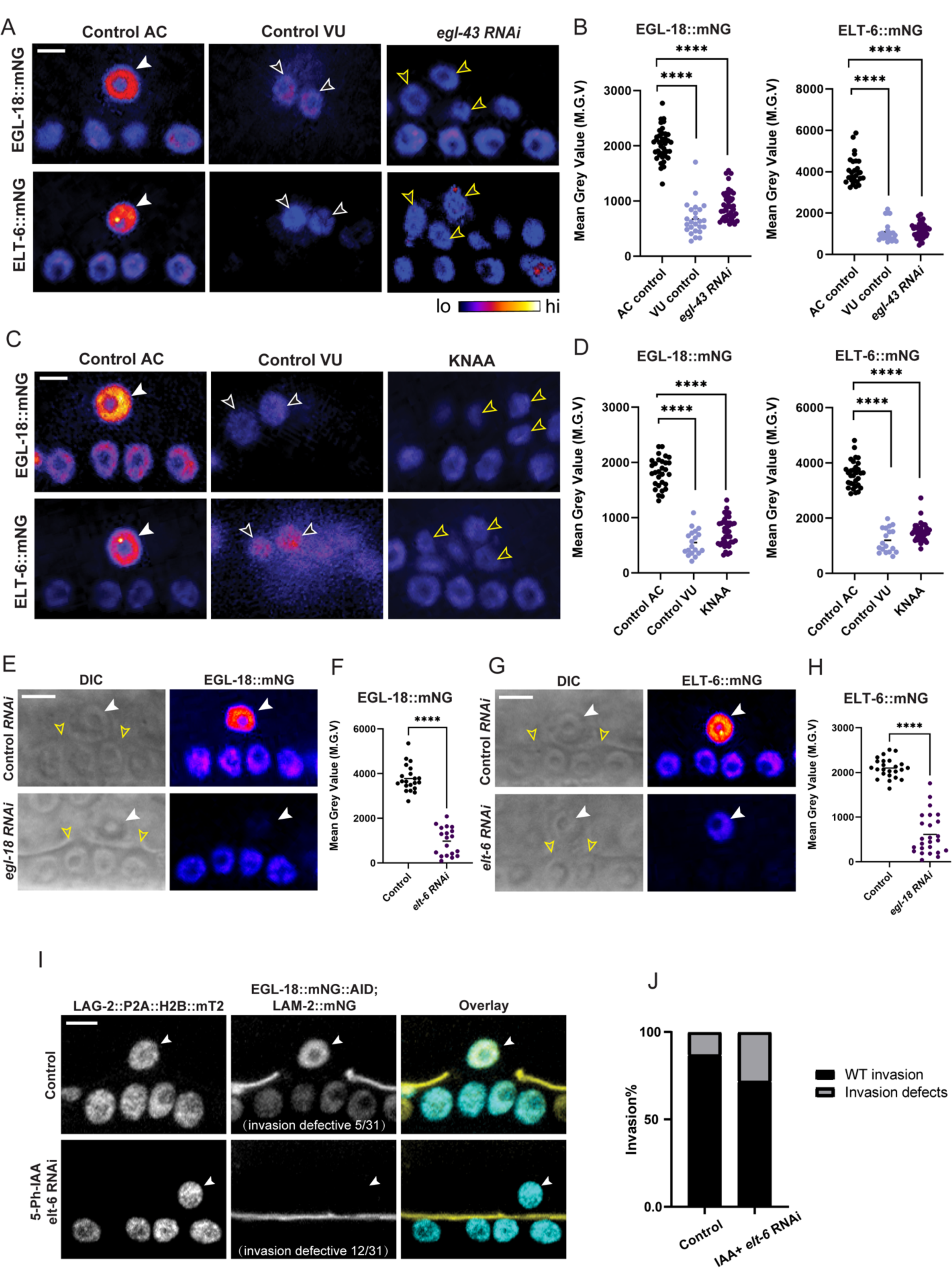
EGL-18 and ELT-6 function downstream of EGL-43 to control anchor cell invasion. A) EGL-43 promotes expression of *elt-6* and *egl-18*. Representative images of the AC in animals expressing EGL-18::mNG(*egl-18(dev210)*) or ELT-6::mNG(*elt-6(dev188)*). Signal intensity is represented using a Fire Lookup table, where blue denotes low expression and red/yellow denotes high expression. EGL-18 (top row) and ELT-6 (bottom row) are expressed highly in the AC (solid white arrows) and lowly in the ventral uterine tissue (outlined white arrows). RNAi knockdown of *egl-43* (yellow arrows) results in loss of EGL-18 and ELT-6 signal in the AC. B) Quantification of EGL-18 and ELT-6 intensity in the AC of animals shown in A). C) Confocal micrographs of EGL-18 and ELT-6 in animals expressing *wrdSi22; egl-43(wy1514)*. Animals were treated in the presence of 0mM or 4mM K-NAA to degrade EGL-43::TagRFP-T::AID* (yellow arrows). D) Quantification of EGL-18 and ELT-6 intensity of the animals shown in C). E) *egl-18* RNAi reduces EGL-18::mNG expression. Left: DIC images of the AC. The solid white arrow indicates the AC; yellow arrows indicate basement membrane where the AC has successfully invaded. Right: fluorescent micrographs of EGL-18::mNG. White arrow indicates the AC. F) Quantification of EGL-18::mNG intensity of the AC from images shown in E. G) *elt-6* RNAi reduces ELT-6::mNG expression. Left: DIC images of the AC. Labelling scheme is the same as in E). Right: fluorescent micrographs of ELT-6::mNG. H) Quantification of ELT-6::mNG intensity of the AC from images shown in G). I) Double depletion of ELT-6 and EGL-18 is sufficient to induce AC invasion defects. Top row: treatment with an empty RNAi vector does not produce a significant invasion phenotype. Bottom row: double treatment with both auxin and *elt-6* RNAi leads to a strong AC invasion phenotype. LAG-2::P2A::H2B::mT2 is an endogenous transcriptional reporter and was used to label the AC and primary VPCs; LAM-2::mNG labels the basement membrane to detect AC invasion. J) Quantification of the animals shown in E). The control condition (0mM 5-Ph-IAA Auxin and empty vector RNAi) yields 87% successful invasion; the double depletion condition (1mM 5-Ph-IAA auxin and *elt-6* RNAi) yields 61% successful invasion). Scale bar = 5μm.

Next, we wanted to test the individual roles of EGL-18 and ELT-6 during AC invasion. We performed RNAi knockdown of *egl-18* and *elt-6* and found that either treatment led to no AC invasion phenotypes (0% defective, n > 30), despite strong knockdown efficiency as measured by endogenous mNG expression (Figure 5E-H). We next wondered if the two TFs may be functioning redundantly due to their overlapping expression pattern. We generated an endogenously tagged mNG::AID allele of EGL-18 and performed double depletion assays by pairing auxin-induced degradation of EGL-18 with RNAi knockdown of *elt-6*. In the absence of auxin, we found that our control RNAi vector induced a weak AC invasion phenotype (13% defective, 87% invasion). However, when we performed both auxin-induced degradation of EGL-18 and *elt-6* RNAi, we observed a strong invasion defect which was similar to what has been reported for EGL-43 RNAi (MEDWIG-KINNEY *et al*. 2020) (39% defective, 61% invasion) (Figure 5E-F). Through these experiments, we have identified two TFs downstream of EGL-43 that possess overlapping expression patterns, and likely function together redundantly to elicit the AC invasion program. This result mirrors prior observations in the epidermis, where *egl-18* functions redundantly with *elt-6* to control seam cell fate (GORREPATI *et al*. 2013).

## Discussion

Here, we demonstrate successful adaptation of the NanoDam system into *C. elegans*, which will enable the *C. elegans* community to perform low-input, tissue-specific profiling using endogenously tagged TFs. First, we show that C- and N-terminal Dam fusions to a GFP nanobody perform well, and that NanoDam generates binding profiles which are comparable to those of TaDa. Second, we demonstrate the utility of NanoDam by profiling different tissues utilizing the same endogenously tagged transcription factor, EGL-43, to demonstrate additional profiling resolution at the single-cell level compared to ChIP-seq and shared negative autoregulation of EGL-43 expression in neurons and ACs. Last, we uncover through NanoDam the GATA factors ELT-6 and EGL-18 as novel players in AC invasion. This finding is significant because despite years of studying AC biology, only a handful of TFs were previously known that regulate AC invasion (*egl-43, hlh-2, nhr-*67 and *fos-*1) (SHERWOOD *et al*. 2005; FERNANDES AND STERNBERG 2007; MATUS *et al*. 2010; SCHINDLER AND SHERWOOD 2011).

### NanoDam vs. Targeted DamID

While previously performing TaDa experiments, we occasionally experienced difficulty in amplifying the GATC-methylated DNA, which led us to incorporate additional PCR cycles in the wet lab protocol (KATSANOS *et al*. 2021). We encountered similar difficulty amplifying DNA in the case of RNT-1 TaDa samples, for which 4 additional PCR cycles had to be performed to obtain sufficient methylation for further analysis. However, more intensive amplification can result in somewhat reduced library complexity due to preferential amplification of a few abundant methylated fragments (MARSHALL *et al*. 2016). Indeed, the RNT-1 TaDa dataset led to identification of substantially fewer statistically significant peaks than NanoDam, therefore both limited methylation and restricted target identification were identified as problems in this case. We reasoned that direct fusions between RNT-1 and Dam could potentially obstruct effective methylation of DNA as a result of fusion protein domain topology, or RNT-1-Dam fusions may be unable to effectively compete with endogenous RNT-1 binding to its targets. NanoDam is less likely to be affected by such limitations since it utilizes endogenously tagged RNT-1 and binding by the nanobody to RNT-1 that already occupies target sequences. It is of note that some variation observed between C-vs N-terminal NanoDam fusions may also be caused by subtle steric interactions that influence the ability of the fusions to methylate proximal DNA.

One key advantage of NanoDam over Targeted DamID (TaDa) is its portability. Using a single NanoDam driver line, users can perform parallel profiling experiments of multiple TFs through a single cross. A second advantage of NanoDam over TaDa is that NanoDam allows for simultaneous profiling of multiple isoforms of TFs depending on the position of the GFP tag (ie. C-terminal vs. N-terminal endogenous labelling). In TaDa, a single isoform is fused directly to Dam methylase, thus restricting the profiling scope. Moreover, due to the inherent design of NanoDam experiments, the technique preserves the *cis* regulatory elements and subsequent endogenous expression of native TFs. This is particularly advantageous if the TF acts in a dose-dependent manner or is natively expressed at low levels to avoid potential toxicity and artificial binding, which may not be representative of the TF’s *in vivo* biology. Finally, another advantage of NanoDam is that it provides greater control of spatial resolution, since target identification occurs at the intersection between the expression domain of the TF and that defined by the promoter of the NanoDam transgene.

We anticipate this tool will be highly valuable to *C. elegans* researchers as the community is continuously producing GFP-tagged TFs at their endogenous loci. A standard ChIP-seq experiment in *C. elegans* requires a large input of approximately 1-2M worms (ASKJAER *et al*. 2014). Like TaDa, NanoDam requires 100-fold lower input to successfully perform TF profiling (KATSANOS *et al*. 2021). In this study, we were able to successfully generate AC-specific NanoDam profiles from gDNA harvested from as few as 10,000 animals. The AC represents ≈0.1% of the total somatic cells in *C. elegans*, providing the most restrictive example to demonstrate the profiling power and specificity of NanoDam.

Taken together, we present NanoDam as a complementary low-input genomics technique which provides resolution, portability, and ease to generate TF-binding profiles in *C. elegans.* With the recent improvements in tissue- and cell-type specific targeted degradation capabilities (XIAO *et al*. 2023), one could simultaneously deplete a protein of interest and query changes in TF-binding genome-wide at single-cell resolution. We are hopeful that this technique will further aid researchers unlock the dissection of tissue and cell-type specific gene regulatory networks that govern many aspects of *C. elegans* biology.

## Acknowledgements

We thank Nuria Flames, Marie Killeen, Kota Mizumoto, Peri Kurshan and members of the Barkoulas, Matus and Shen labs for critical reading of this manuscript. Some strains were provided by the *Caenorhabditis* Genetics Center (CGC), funded by the NIH Office of Research Infrastructure Programs (P40 OD010440).

## Funding

We acknowledge the support from the European Commission (ROBUSTNET-639485). CY was supported by the Human Frontiers Science Program (LT000127/2016-L). DK was the recipient of an Imperial College President’s PhD scholarship. DQM and TNM-K were funded by the National Institutes of Health (R01GM121597 and F31HD100091, respectively). KS is a Howard Hughes Medical Institute Investigator.

### Data Availability

Sequencing data has been deposited in Gene Expression Omnibus (GEO) with Accession numbers: GSE227552, GSE232946.

### Contact for Reagent and Resource Sharing

Further information and requests for reagents and resources should be directed to and will be fulfilled by the lead contacts, Callista Yee and Michalis Barkoulas.

## Supplemental Material

**Table S1: List of peaks from RNT-1 TaDa, RNT-1 NanoDam and RNT-1 ChIP-seq analyses**

**Table S2: List of peaks from EGL-43 ChIP-seq, EGL-43 AC and DA NanoDam**

**Table S3: Gene Ontology analysis of peaks from EGL-43 ChIP-seq and NanoDam experiments**

**Table S4: Tissue enrichment analysis of peaks from EGL-43 NanoDam experiments**

**Table S5: Strains used in this study**

**Table S6: Primers used in this study**

---

**Figure S1:**
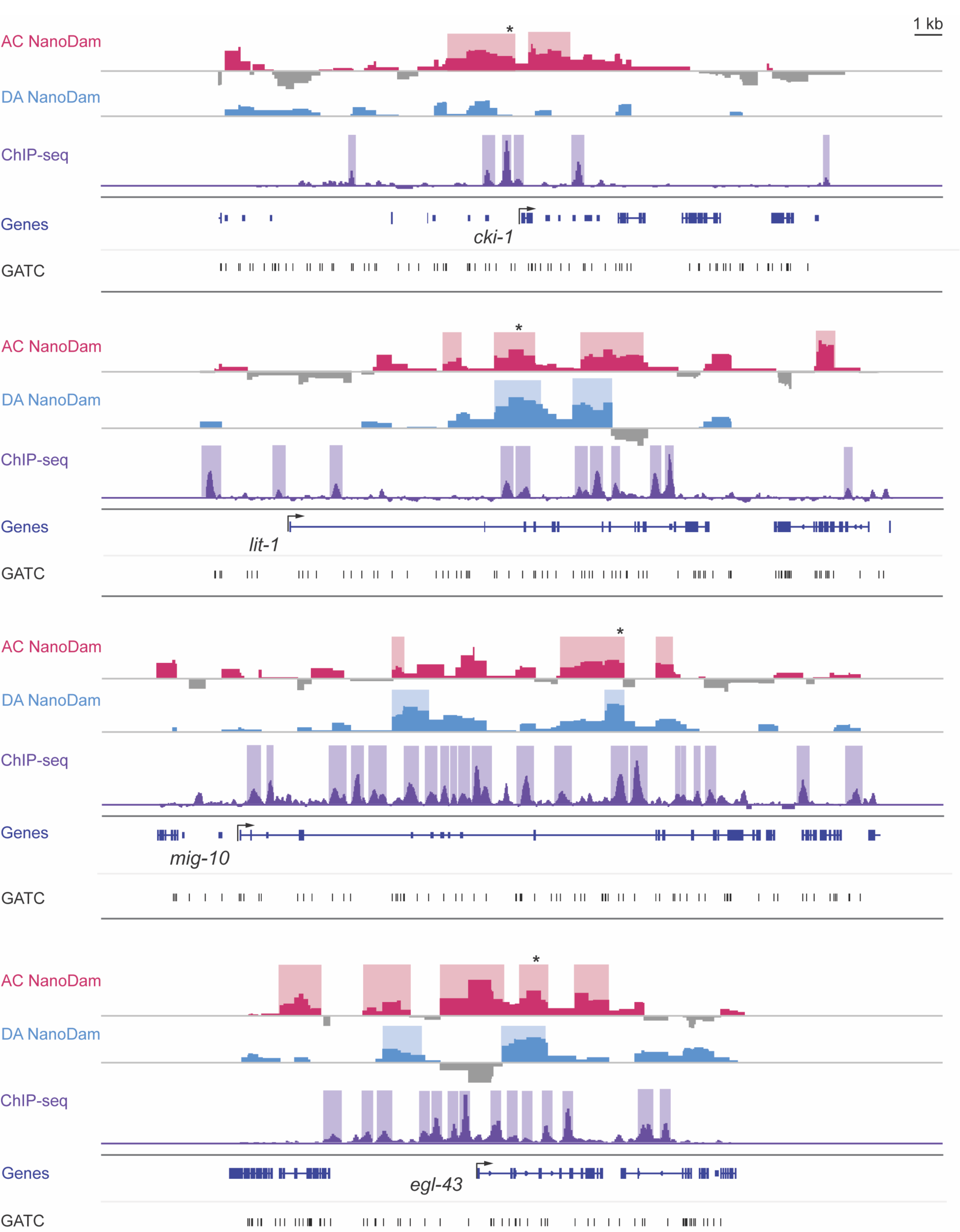
EGL-43 NanoDam and ChIP-seq binding profiles at loci of genes that function genetically downstream of *egl-43*. For each locus, binding profiles for EGL-43 AC NanoDam (red), EGL-43 DA NanoDam (light blue) and EGL-43 ChIP-seq (purple) are presented relative to genomic position (dark blue) and GATC sites. Computed peaks are denoted by lightly shaded rectangles on the NanoDam and ChIP-seq tracks. Genetic loci for *cki-1, lit-1, mig-10* and *egl-43* are shown respectively. Asterisks indicate regions overlapping with NanoDam and ChIP-seq peaks that were tested for binding using ChIP-qPCR.

